# Randomized Spatial Barcoding for Time-Lapse Flow Cytometry

**DOI:** 10.64898/2026.07.20.738363

**Authors:** Xiaoyao Chen, Masashi Ugawa, Sadao Ota

## Abstract

Tracking suspended cells over multiple time points at the single-cell level remains challenging because existing flow-based methods cannot preserve cell identity while maintaining high throughput. Here, we present RASPBerry, a hydrogel-based spatial barcoding platform for time-lapse flow cytometry. RASPBerry generates unique barcodes by randomly co-encapsulating fluorescent beads with individual cells in hydrogel droplets, eliminating the need for predefined barcode patterns or specialized optical instrumentation. We integrate RASPBerry with acoustofluidic imaging flow cytometry to enable time-lapse imaging flow cytometry of suspended cells. The platform identifies more than 17,000 hydrogel droplets with 99.8% matching accuracy. We further demonstrate time-lapse tracking of more than 10,000 suspended cells and quantify stress-induced nuclear morphological changes in more than 5,000 individual cells. RASPBerry provides a simple, scalable, and broadly accessible strategy for time-lapse imaging flow cytometry, expanding the capability for dynamic single-cell analysis of suspended cells.

## 1. Introduction

The dynamic nature of cellular systems and their responses to environmental stimuli necessitate robust methods for tracking individual cells over time.^[1,2]^ Although single-cell sequencing provides valuable insights through endpoint expression profiles,^[3,4]^ its destructive nature makes optical analysis methods, such as time-lapse microscopy, essential for tracking and monitoring cells in real time.^[2]^ Moreover, recent advances in optical pooled screening have further increased the need for imaging at multiple time points, as genetic barcode readout often relies on sequential imaging of cells over time.^[5–7]^ However, single-cell tracking has largely been limited to adherent cells, which remain relatively stationary. In contrast, tracking suspended cells—which holds significant potential for understanding hematopoietic cell differentiation, immune cell regulation, and stem-cell differentiation for regenerative medicine^[8–10]^—remains challenging. Multi-well systems can be used for this purpose, but they face limitations in scalability and potential effects on cell health due to cellular isolation.^[11,12]^ Genetic barcoding strategies based on color-coded labeling of subcellular structures also have limited scalability.^[13,14]^ Alternative approaches, such as single-cell barcoding systems using microdroplets, can improve scalability but still suffer from restricted medium exchange and limited intercellular interaction.^[15,16]^

Flow-based analysis methods such as flow cytometry and imaging flow cytometry have revolutionized the study of suspended cells by enabling high-throughput analysis and diverse applications.^[17–20]^ However, these methods lack the ability to track individual cells pooled in solution over time, and time-lapse measurements have therefore been limited to the population level. Recently, a single-cell barcoding system for flow cytometry based on microdisks was developed.^[21,22]^ However, this technology requires (i) endocytosis of the microdisks, which limits the cell types that can be used, and (ii) a spectral analyzer equipped with a near-infrared laser source, which is not standard equipment in many laboratories and core facilities. Therefore, despite the high throughput and scalability of this barcoding technology, its practicality remains limited. Hydrogel microparticles with optically defined spatial codes have also been developed for high-throughput biomolecular analysis, demonstrating the utility of encoded hydrogel materials as scalable assay carriers.^[23]^ However, when applied to single-cell time-lapse tracking, predefined optical or lithographic encoding strategies would require generating and assigning distinct patterns to large numbers of cell-containing hydrogels, which can limit practical throughput and implementation.

To address these challenges, we introduce RASPBerry (RAndomized SPatial Barcoding with Encapsulation of Retained paRticles in hYdrogels), a hydrogel-based spatial barcoding platform for time-lapse flow cytometry of suspended cells. RASPBerry encodes each cell-containing hydrogel droplet by the randomized positions of retained fluorescent beads, producing a spatial barcode that can be read using only a single fluorescence channel on a standard fluorescence microscope. Unlike approaches that require intracellular uptake of barcode materials, specialized spectral instrumentation, or predefined optical patterns for individual hydrogel particles, RASPBerry generates barcode diversity stochastically during cell and bead co-encapsulation, enabling scalable implementation with minimal optical complexity. Because RASPBerry requires only co-encapsulation of cells with barcoding beads, it enables single-cell time lapse imaging at an unprecedented scale of over 10,000 cells. Furthermore, we demonstrated that it can track and analyze morphological changes of immune cell lines induced by stress for more than 5,000 cells.

## 2. Results

### 2.1 Concept of the randomized spatial barcoding system

The key idea of our cell barcoding system, RASPBerry, is to encapsulate fluorescent beads together with a cell in each droplet and use the relative spatial positions of the beads as a unique barcode for that droplet. To achieve this, two requirements must be satisfied: the positions of the beads must be preserved throughout the measurement, and the bead configuration must differ among droplets. The first requirement is met by using hydrogel droplets, in which the beads are immobilized within the hydrogel matrix. The second requirement is met by encapsulating multiple beads that are small relative to the droplet size, thereby generating a large number of possible spatial arrangements. The number of unique barcodes can be estimated from the number of possible bead configurations within a droplet. For example, in a spherical droplet with a diameter of 50 µm containing beads with a diameter of 5 µm, more than 10^10^ unique spatial configurations can be generated even with only four beads per droplet and a spatial resolution of 2 µm (Supplementary Information). However, reading and matching spatial barcodes in spherical droplets is computationally demanding because of their high rotational symmetry.

To implement hydrogel droplets suitable for flow-based barcode imaging, we took advantage of the shape-preserving property of hydrogel droplets and designed obround, or stadium-shaped, droplets (Figure 2a). This geometry is advantageous for time-lapse imaging flow cytometry for three reasons. First, it facilitates in-focus imaging of both the beads and the cell within the droplet. In our flow system, the droplets are acoustically focused such that the flat plane of each droplet is aligned parallel to the imaging plane (Figure 2b). By designing the droplet thickness to be smaller than the depth of focus, both the beads and the cell can be imaged in focus. Second, the obround geometry provides a consistent droplet orientation with respect to the imaging plane under acoustic focusing (Figure 2b). Third, elongation along one axis breaks the rotational symmetry of the droplet in the two-dimensional image, thereby reducing the computational complexity of image matching (Figure 2c).

**Figure 1.**
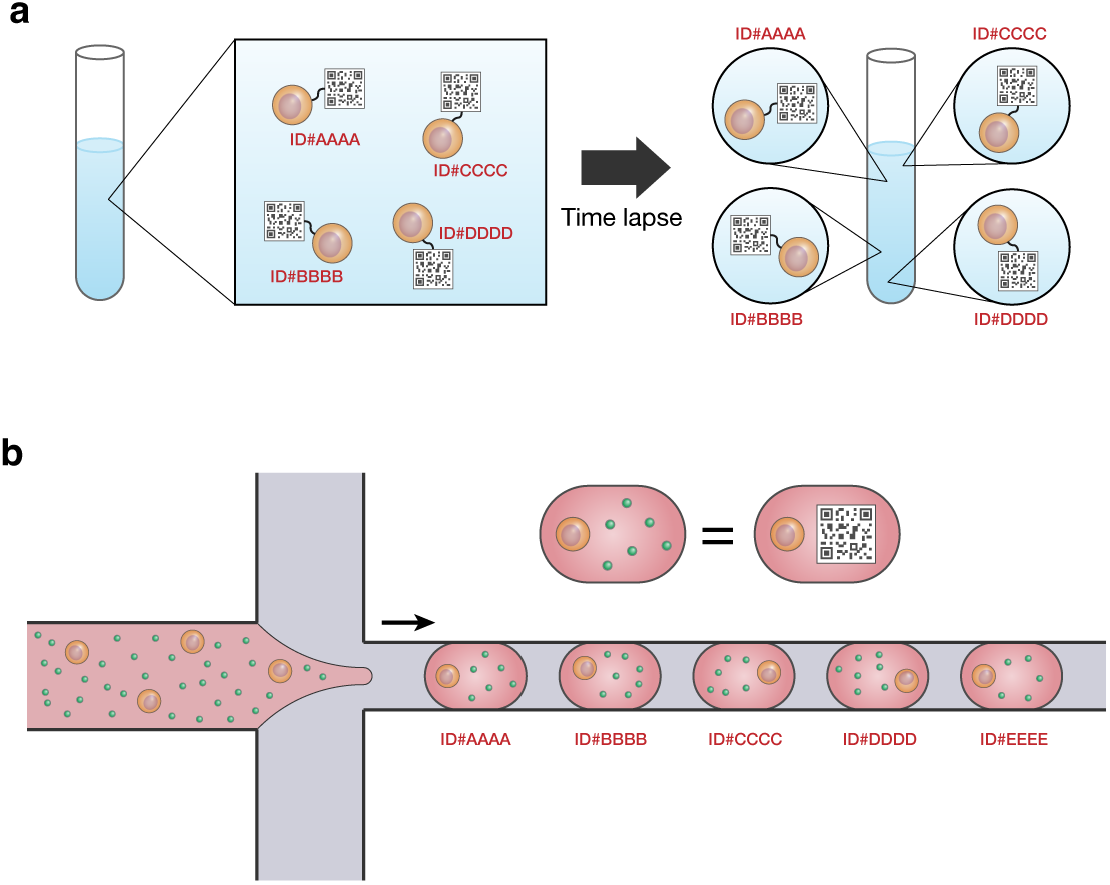
Concept of the RASPBerry system. (a) Individual suspended cells can be identified by attaching a spatial barcode that can be read out by imaging. This enables each cell to be tracked over time while in suspension. (b) Spatial barcodes are generated by randomly encapsulating fluorescent beads within hydrogel droplets. The random positions of the beads serve as a spatial barcode. These barcodes are associated with individual cells by co-encapsulating cells and beads within the same hydrogel droplets.

**Figure 2.**
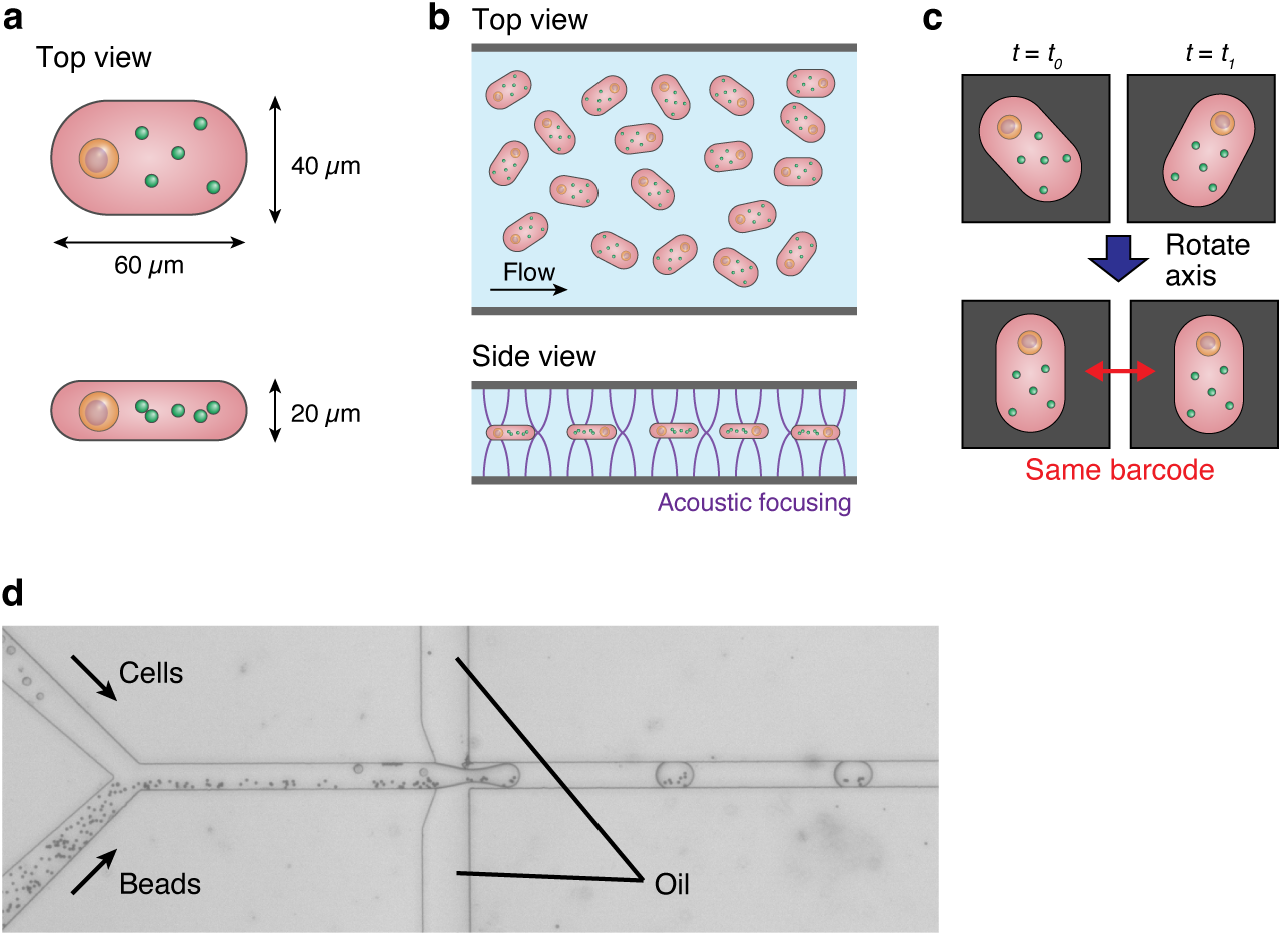
Design of RASPBerry droplets. (a) Geometry of RASPBerry droplets. Viewed from the top, the droplets have an obround or stadium shape. (b) Orientation of RASPBerry droplets in a 1D acoustofluidic focusing channel. Acoustic standing waves generated in the vertical direction will focus the droplets at the center while simultaneously orienting their flat face perpendicular to the direction of the standing waves. (c) The obround shape enables identification of the same barcode by aligning the major axis of the droplet. (d) Microscopic image of the droplet generation device. Cell and bead suspensions in alginate-EDTA PBS solution are combined and subsequently passed through the droplet generating junction. The oil phase contains 1% acetic acid that allows the in-device gelation of the alginate.

### 2.2 Fabrication and validation of barcoded hydrogel droplets

Hydrogel droplets were fabricated using a custom microfluidic droplet generator. We chose an alginate gel system similar to that used in previous reports.^[24,25]^ Alginate gels are suitable for our barcoding system because they are cell-compatible,^[26,27]^ have a refractive index close to that of water to minimize image distortion,^[27,28]^ and can be gelled rapidly to preserve the shape of hydrogel droplets.^[25,29]^ Suspensions of cells and beads were prepared separately in alginate-EDTA PBS solution (see Methods for formulation). The two suspensions were combined in the microfluidic channel and subsequently passed through the droplet-generation junction, where droplets were formed in an oil phase containing 0.1% acetic acid (Figure 2d). The droplets then passed through an 18-mm-long channel with a cross-sectional dimension of 20 × 40 µm (Supplementary Figure 1a), allowing the alginate droplets to gel while being confined by the channel walls and thereby forming obround hydrogel droplets. The droplet length was adjusted to approximately 60 µm, as longer droplets tended to fold after gelation (Supplementary Figure 1, b and c).

As a proof of concept for the barcoding system, we first evaluated the identification of RASPBerry droplets deposited in a well. To enable high-throughput imaging of the droplets, we used a line-scan imaging setup consisting of multiple continuous-wave lasers and a high-speed sCMOS camera (Supplementary Figure 2). Two-dimensional images of hydrogels deposited in a 10 × 10 mm cover-glass-bottom well were acquired by line illumination while translating the sample stage in the direction orthogonal to the illumination axis (Figure 3, a and b). The well was then rotated by 90° without disturbing the relative positions of the deposited droplets, and a second image dataset was acquired using the same procedure (Supplementary Figure 3a). From the raw images, individual hydrogel droplets were segmented to generate a dataset containing both the bead-based barcode patterns and the images of cells within the hydrogel droplets (Figure 3c). Because the relative positions of the hydrogel droplets within the well were preserved during rotation, each droplet image acquired before rotation could be paired with the corresponding image acquired after rotation, providing ground-truth droplet identities. Using the bead-based barcode patterns, we then applied a pattern-matching algorithm to identify hydrogel droplets based on the bead positions within each droplet (Supplementary Information). Among a total of 17,018 RASPBerry droplet images obtained from 4 wells, identical droplets were identified with an accuracy of 99.8%, with only 35 mismatches, most of which occurred in droplets containing two or fewer beads (Supplementary Figure 3c). These results demonstrate that RASPBerry enables large-scale single-cell identification with high accuracy.

**Figure 3.**
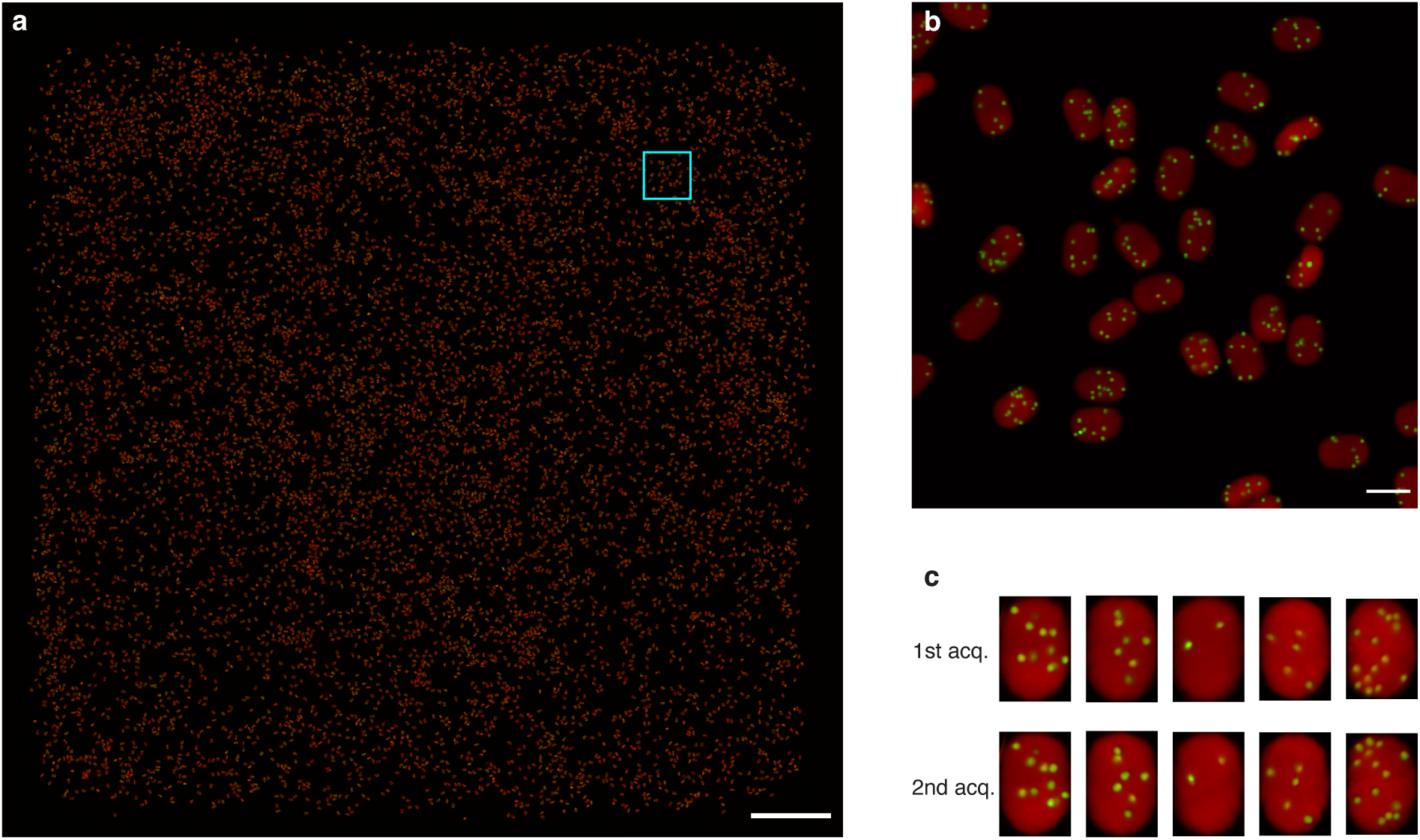
Imaging of RASPBerry droplets in a well. (a) Full view of a single well containing RASPBerry droplets. Red: alginate gel; green: beads. Scale bar = 1 mm. (b) Magnified view of the region outlined by the cyan box in (a). Scale bar = 50 µm. (c) Rotated images of individual RASPBerry droplets from two separate acquisitions. Each pair of images in a column corresponds to the same droplet.

### 2.3 Time-lapse flow cytometry with RASPBerry droplets

To flow the RASPBerry droplets for 2D time-lapse imaging flow cytometry, we developed a 1D acoustofluidic focusing microfluidic device similar to that previously described (Figure 4a).^[30,31]^ The advantage of this device is that (i) it orients the obround-shaped hydrogel droplets parallel to the imaging plane and (ii) focuses the hydrogel droplets at the imaging plane even at slow flow velocities that are required for imaging. Here, we used a glass-PDMS-glass device which allows acoustic standing waves to form only in the vertical direction of the channel (Figure 4b). Furthermore, we introduce a weak sheath flow to keep the droplets away from the side walls where the flow velocity drops compared to other positions (Figure 4c). Using this design, we confirmed that the RASPBerry droplets flowed with a consistent orientation and uniform flow velocity (Figure 4, d and e). These characteristics allow both accurate cell imaging and barcode readout necessary for time-lapse imaging flow cytometry.

**Figure 4.**
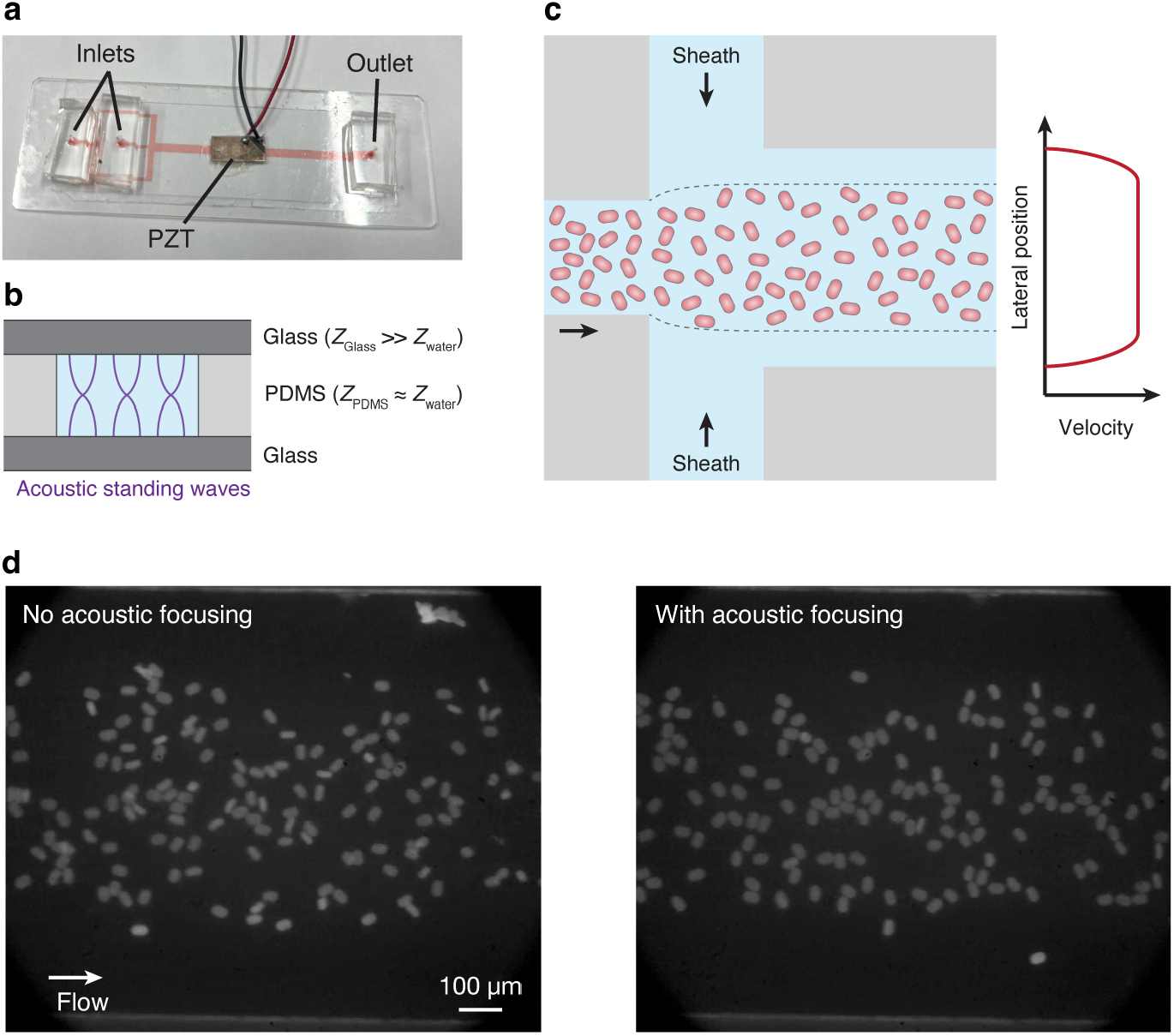
1D acoustofluidic device for 2D imaging flow cytometry with RASPBerry droplets. (a) Photo of the 1D acoustofluidic device. The device consists of with two glass slides with a PDMS layer in between. Additional PDMS blocks are attached at the inlets and outlet for tubing connections. A piezoelectric transducer (PZT) is attached to the glass above the channel. (b) Side-view schematic of the channel. Acoustic standing waves are generated in the vertical direction due to the high reflectivity at the interface between water and glass. In contrast, standing waves are not generated in the horizontal direction due to the low reflectivity at the interface between water and PDMS. (c) Top view schematic of the channel. Weak sheath flows on both sides of the channel keep the RASPBerry droplets away from the channel walls, allowing them to flow at uniform velocity. (d) Images of RASPBerry droplets flowing through the channel without acoustofluidic focusing (left) and with acoustofluidic focusing (right). Without focusing, the droplets are randomly oriented, whereas with focusing, the droplets are oriented with their flat faces parallel to the imaging plane.

To investigate the feasibility of RASPBerry for time-lapse flow cytometry, we performed time-lapse tracking of cells encapsulated in the hydrogel droplets. To maintain high cell viability during encapsulation, we first modified the droplet generation process to use cell-compatible conditions. The cells were suspended in alginate-GEDTA PBS solution prior to loading into the hydrogel-forming device. Replacing EDTA with GEDTA reduced the acidity required for alginate gelation,^[32]^ thereby improving cell viability. Furthermore, the droplets were collected directly into culture medium to neutralize the buffer as soon as gelation was complete. As a result, cell viability after encapsulation was maintained at 90%. Maintaining cell viability is challenging when individual cells are isolated in separate wells of a multiwell plate because they are cultured at a very low cell density.^[11]^ Together, these results demonstrate that RASPBerry maintains sufficient cell viability for live-cell imaging and time-lapse analysis of suspended single cells.

Using this hydrogel encapsulation strategy, we performed the single-cell analysis of cells treated with hydrogen peroxide to induce oxidative stress. The cells were first imaged with our custom 2D imaging flow cytometry system after encapsulation into the RASPBerry droplets (Figure 5a). Then the cells were treated with hydrogen peroxide for 4 hours while inside the droplets and were subsequently imaged again. A total of 54,619 and 45,057 droplets were imaged before and after treatment, respectively, and 10,926 droplets containing cells were correlated. From these droplets, images of 5,564 cells without occlusion and multiplets were extracted and the area of the nucleus for each cell in each acquisition was measured (Figure 5, b and c). From this measurement, we observed that the relative size change showed a normal distribution with a mean and standard deviation of –7.7 ± 12.7% (Figure 5d). Furthermore, with the ability to track and image each cell at the single cell level, we were able to extract the shape changes of the nuclei which had a significant size change (Figure 5, e and f). This analysis demonstrates the ability to capture rare shape changes of suspended cells within a large population.

**Figure 5.**
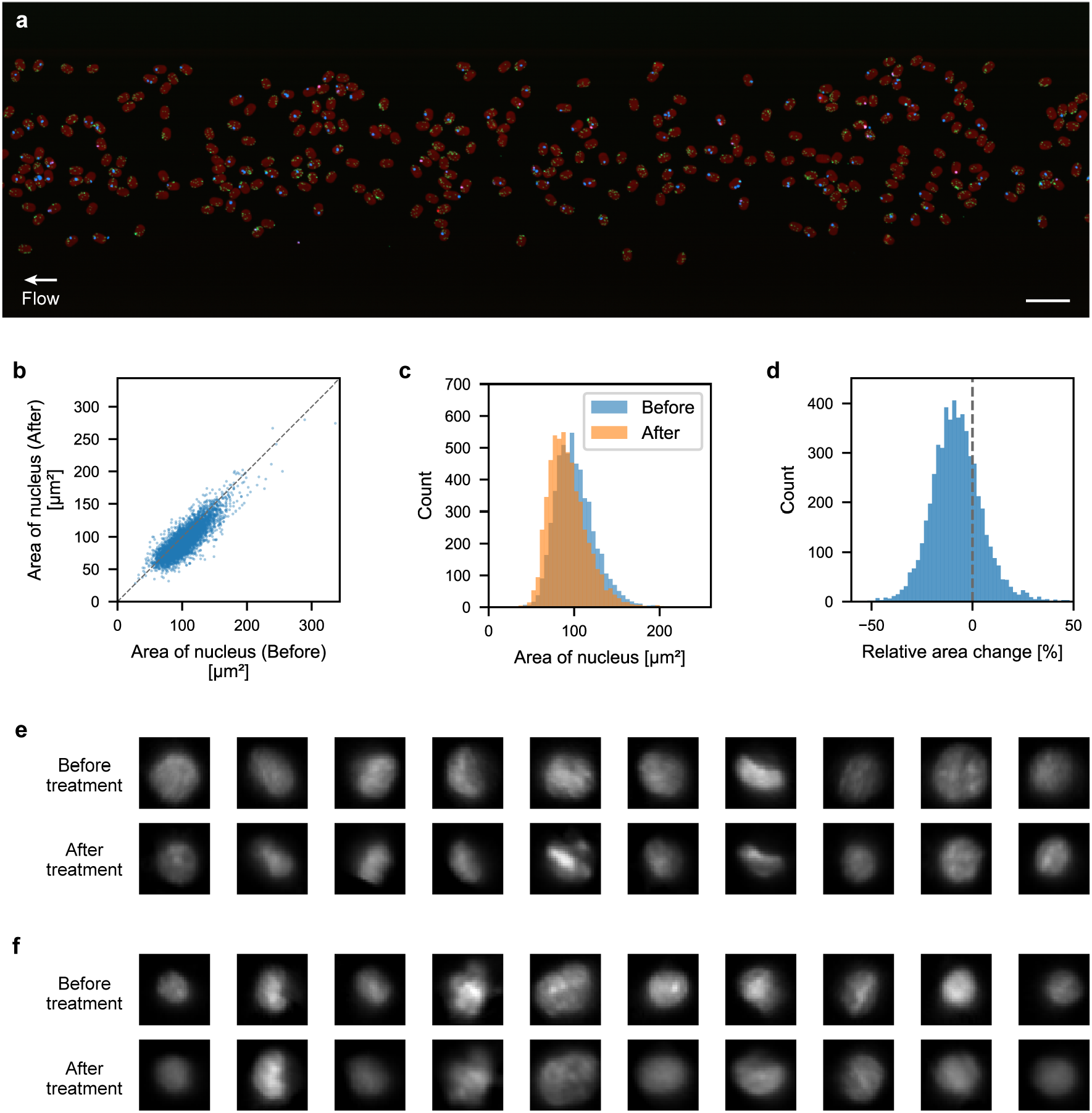
Time-lapse imaging of Jurkat cells treated with hydrogen peroxide. (a) Reconstructed image from a 2D imaging flow cytometry acquisition. Red: alginate gel; green: beads; cyan: cell nucleus. Scale bar = 200 µm. (b, c) Scatter plot (b) and histograms (c) of nucleus area, before and after treatment obtained from 5,564 cell images. (d) Relative change in the area of the nucleus following treatment. (e, f) Example images of nuclei which decrease (e) and increase (f) in area following treatment.

## 3. Discussion

The main advantage of our bead-based barcoding system is the scalability. Because it utilizes the spatial positioning of multiple beads, it can easily scale to over thousands of unique barcodes with only a single color channel. The number of unique barcodes it can create can be roughly estimated by the number of spatial positions the beads can take inside the hydrogel droplet, and even with only 4 beads per droplet, this already corresponds to 2.7 × 10^8^ barcodes (Supplementary Information), which is far more than we can acquire. Given that the number of beads per droplet will vary, the actual number of barcodes that can be created will be much larger.

Instead, the limitation of scalability comes from the number of cells that can be imaged in a certain amount of time. Because the RASPBerry droplets occupy a larger space than cells, it reduces the amount of cells that can be imaged in the same amount of time. This can be estimated to be 200 droplets/sec at the maximum with optimal resolution (Supplementary Information). Furthermore, not all droplets will contain cells due to the Poisson distribution during encapsulation.^[33]^ This can be tuned based on the requirements of the experiment: whether having multiple cells in a droplet is acceptable and the recovery rate of the analysis. For a high recovery rate, the parameter lambda for Poisson distribution should be set as 0.1, and the droplets containing cells will be around 10% and most of them will contain only one. If data dropout is acceptable, the lambda can be set as 1.0 and one-third of the droplets will contain one cell and one-third will contain multiple cells. Therefore, the number of cells that can be imaged will be reduced by a factor of 3–10 compared to the number of droplets that can be imaged.

A key factor to performing accurate time-lapse measurements is the barcode matching error. In the demonstration we performed in a well, we were able to achieve a high accuracy due to the completeness of the dataset in each round (i.e. all barcodes had a corresponding pair). However, in actual flow cytometry measurements, hydrogel droplets can be lost between measurements, misoriented during imaging, or misanalyzed during image processing, and therefore, the corresponding pair may not exist during matching. Even if such dropouts of hydrogel droplets are a small fraction, it may significantly affect the accuracy of matching. The error in the matching will result in the dropout of data, and this will become larger when increasing the measurement timepoints.

The resolution of our system is both scalable and limited depending on multiple factors. One factor to take into account is the tradeoff between resolution and throughput. The higher the resolution, the field-of-view becomes smaller, resulting in a lower throughput. To overcome this, the droplets and barcode beads can be made smaller. However, smaller droplets are difficult to create with the microfluidic droplet generation system we used. Also, the thickness of the droplets, which needs to be the shortest dimension of the hydrogel droplet, cannot be made smaller than the cell size. Even when compromising the throughput, increasing the resolution will be difficult because it will require a stricter focus of the cells due to a smaller depth of focus. In the current system, the vertical position of the cells and the beads can deviate inside the hydrogel, and that will result in an inconsistent focus during imaging at higher resolution. A couple of ways to solve this are to use extended depth of focus methods^[34,35]^ or oblique-plane microscopy^[30,31,36,37]^ to capture multiple depths at once.

For live-cell applications, cell compatibility with alginate gel is essential. Many cell types have been shown to remain viable in alginate.^[25–27]^ However, during our experiments, we observed that although the cells remained viable, their ability to proliferate appeared to be reduced. Furthermore, this ability to proliferate appeared to depend on the alginate concentration. Whether this is a chemical effect or a mechanical effect is uncertain. Nevertheless, this limited proliferation is advantageous for our application because cell proliferation within the hydrogel can distort both the hydrogel and the embedded barcode, resulting in matching errors. Developing barcoding technologies that enable cell tracking during proliferation will be an important direction for future research.

Future expansion of this technology can be directed in multiple ways. One direction is to expand the dimension of the imaging to perform 3D time-lapse imaging. Another direction is to observe cell secretion, such as metabolites and extracellular vesicles, while maintaining the intercellular interaction. Furthermore, time-lapse capability can be used to combine multiple modalities that are measured at different timepoints. This can be useful for performing multi-parameter perturbation screening in applications such as optical pooled screening and directed evolution.

## 4. Conclusion

In conclusion, we have developed a bead-position-based barcoding system using hydrogel droplets which is a versatile and scalable approach for tracking and analyzing suspended cells in time-lapse flow cytometry. We have successfully demonstrated time-lapse imaging of more than 10,000 cells at the single-cell level which was difficult with conventional high-throughput well-based platforms or imaging flow cytometry systems. The simplicity and adaptability of this method make it a promising tool for studying cellular dynamics in various contexts, paving the way for future advances in our understanding of cellular behavior and responses to environmental stimuli.

## 5. Methods

### Microfluidic device fabrication

For the RASPBerry droplet fabrication, a polydimethylsiloxane (PDMS) microfluidic device was used. The channel patterns on the PDMS were created using a master mold fabricated by standard soft lithography with SU-8 photoresist (Nippon Kayaku) on a silicon wafer. The patterned PDMS was attached to a glass slide using a plasma cleaner (Harrick Plasma). The glass-PDMS-glass device used for the acoustofluidic focusing was fabricated by bonding a printed PDMS layer with a thickness of 100 µm (CSTEC) to glass slides (Matsunami Glass) with the plasma cleaner.

### Line-scan microscopy setup

Three continuous-wave lasers with wavelengths of 405, 488, 637 nm were combined using dichroic mirrors (Chroma). The combined beam was shaped to a line beam using a combination of cylindrical lenses (Thorlabs) and was directed to a 10× objective lens with an NA of 0.3 (UPLFLN10×2, Olympus). The fluorescence was separated from the excitation with a multiband dichroic mirror (ZT405/488/635rpc-UF1, Chroma) and was imaged on to a sCMOS camera (Zyla 5.5, Andor) using a tube lens (AC508-200-A-ML, Thorlabs). The sCMOS camera was equipped with a multibandpass filter (ZET405/488/561/640mv2, Chroma). The lasers and camera were operated by an external trigger signal generated by a combination of a function generator (Rigol) and an Arduino microcontroller. The camera was operated at 2 kHz while the lasers were operated alternatingly and in sync with the camera. Line images were acquired using Solis (Andor) and were reconstructed into a 2D image with a custom code written in Python. Details of the setup are depicted in Supplementary Figure 2.

### Fabrication of RASPBerry droplets

Fluorescent beads with a diameter of 4.5 µm (Polysciences) were suspended in a solution of 1% alginate, 50 mM EDTA in PBS. For the cell encapsulation, a solution of 1% sodium alginate, 50 mM GEDTA in PBS was used for both beads and cell suspension. For the oil phase, Droplet Generation Oil for EvaGreen (BioRad) with 0.1% of acetic acid was used. The aqueous phase was driven with a pressure pump (Fluigent), and the oil phase was driven with a syringe pump (Harvard Apparatus). The oil phase was fixed at a flow rate of 2.3 µL while the pressure of the aqueous phase was adjusted around 1,150 mbar by confirming the length of the generated droplets on a microscope. The collected droplets were removed from the oil phase with repeated washing using 20% 1H,1H,2H,2H-perfluoro-1-octanol in HFE7200 and was resuspended in a buffer containing 10 mM Tris-HCl, 137 mM NaCl, 2.7 mM KCl, and 1.8 mM CaCl_2_.

### Identification of RASPBerry droplets imaged in wells

Wells in a cover glass chamber with 10 × 10 mm well (IWAKI) was pre-treated with poly-L-lysine. A suspension of the RASPBerry droplets were added to the treated wells, and the droplets were allowed to deposit to the bottom. Using the line-scan microscope and two orthogonally oriented motorized stages (Optosigma), the whole area was scanned for each of the wells. After the first acquisition, the chamber was rotated 90 degrees and the second acquisition was performed (Supplementary Figure 3a). The acquired images were stitched based on the overlap between each line-scanned image. The relative position between the two acquisitions were adjusted based on the overall position of the imaged droplets. Identical droplets were identified based on the distance between the position in the first and second acquisitions. 17,018 images of droplets with the flat face parallel to the imaging plane were extracted based on the area and fluorescence intensity (Supplementary Figure 3b). The long axis of the droplets were aligned by principal component analysis as previously described.^[30]^ Here, the relative angle between the first and second acquisitions were not used, and the images from each acquisition were processed independently. The identification of identical droplets were performed using a template matching algorithm in Open CV. After the correlation score for all droplet combinations were calculated, an image pair was considered a correct match only when within all the images from the second acquisition, the image having the best correlation score with the image from the first acquisition was the corresponding pair and vice versa.

### Operation of acoustofluidic focusing device

A sine wave generated with a function generator (Rigol) was amplified using a bipolar amplifier (HSA4101, NF Corporation). The amplified signal of 15 Vpp was applied to the piezoelectric transducer attached to the acoustofluidic focusing device. The frequency of the sine wave was tuned based on the focusing of the RASPBerry droplets between the range of 7.5–8.1 MHz. The inner sample flow and sheath flow were each flowed at a flow rate of 2.6 and 1.4 µL/min, respectively, with syringe pumps (Harvard Apparatus).

### Cell culture

Jurkat cells (RCB3052) were provided by the RIKEN BRC through the National Bio-Resource Project of MEXT, Japan. They were cultured in RPMI-1640 with 10% FBS, 1% L-Alanyl-L-glutamine, and 1% of 100× Antibiotic-Antimycotic.

### Time-lapse imaging flow cytometry

The RASPBerry droplets containing cells were suspended in the above culture medium with the concentration of Ca^2+^ adjusted to 1.8 mM by additional CaCl_2_. The cells in the RASPBerry droplets were stained with Hoechst 33342 before loading to the acoustofluidic focusing device. The suspended RASPBerry droplets were loaded in a PTFE tubing connected to a syringe using a syringe pump in withdrawing mode. The tubing was connected to the acoustofluidic focusing device, and the suspension was flowed at 2.6 µL/min. The image was acquired for 30 min and the suspension was collected in a 1.5 mL tube. The collected RASPBerry droplets were resuspended in 1.8 mM Ca^2+^ culture medium with 150 μM hydrogen peroxide, and were incubated for 4 h. After 4 h, the droplets were resuspended in 1.8 mM Ca^2+^ culture medium and re-stained with Hoechst. This was then flowed through the acoustofluidic focusing device similarly to the first acquisition. The second acquisition was performed for 30 min. The images acquired from both acquisitions were processed with a custom code written in Python.

## References

1. Spiller, D. G., Wood, C. D., Rand, D. A., & White, M. R. H. (2010). Measurement of single-cell dynamics. Nature, 465(7299), 736–745. 10.1038/nature09232

2. Skylaki, S., Hilsenbeck, O., & Schroeder, T. (2016). Challenges in long-term imaging and quantification of single-cell dynamics. Nature Biotechnology, 34(11), 1137–1144. 10.1038/nbt.3713

3. Papalexi, E., & Satija, R. (2018). Single-cell RNA sequencing to explore immune cell heterogeneity. Nature Reviews. Immunology, 18(1), 35–45. 10.1038/nri.2017.76

4. Stuart, T., & Satija, R. (2019). Integrative single-cell analysis. Nature Reviews Genetics, 20(5), 257–272. 10.1038/s41576-019-0093-7

5. Feldman, D., Singh, A., Schmid-Burgk, J. L., Carlson, R. J., Mezger, A., Garrity, A. J., Zhang, F., – Blainey, P. C. (2019). Optical Pooled Screens in Human Cells. Cell, 179(3), 787–799.e17. 10.1016/j.cell.2019.09.016

6. Kudo, T., Meireles, A. M., Moncada, R., Chen, Y., Wu, P., Gould, J., Hu, X., Kornfeld, O., Jesudason, R., Foo, C., Höckendorf, B., Corrada Bravo, H., Town, J. P., Wei, R., Rios, A., Chandrasekar, V., Heinlein, M., Chuong, A. S., Cai, S., … Lubeck, E. (2024). Multiplexed, image-based pooled screens in primary cells and tissues with PerturbView. Nature Biotechnology, 1–10. 10.1038/s41587-024-02391-0

7. Gu, J., Iyer, A., Wesley, B., Taglialatela, A., Leuzzi, G., Hangai, S., Decker, A., Gu, R., Klickstein, N., Shuai, Y., Jankovic, K., Parker-Burns, L., Jin, Y., Zhang, J. Y., Hong, J., Niu, X., Costa, J. A., Pezet, M. G., Chou, J., … Gaublomme, J. T. (2024). Mapping multimodal phenotypes to perturbations in cells and tissue with CRISPRmap. Nature Biotechnology, 1–15. 10.1038/s41587-024-02386-x

8. Laurenti, E., & Göttgens, B. (2018). From haematopoietic stem cells to complex differentiation landscapes. Nature, 553(7689), 418–426. 10.1038/nature25022

9. Liston, A., & Gray, D. H. D. (2014). Homeostatic control of regulatory T cell diversity. Nature Reviews Immunology, 14(3), 154–165. 10.1038/nri3605

10. Matsuo-Takasaki, M., Kambayashi, S., Hemmi, Y., Wakabayashi, T., Shimizu, T., An, Y., Ito, H., Takeuchi, K., Ibuki, M., Kawashima, T., Masayasu, R., Suzuki, M., Kawai, Y., Umekage, M., Kato, T. M., Noguchi, M., Nakade, K., Nakamura, Y., Nakaishi, T., … Hayashi, Y. (2024). Complete suspension culture of human induced pluripotent stem cells supplemented with suppressors of spontaneous differentiation. eLife, 12, RP89724. 10.7554/eLife.89724

11. Munoz, A., & Morachis, J. M. (2022). High efficiency sorting and outgrowth for single-cell cloning of mammalian cell lines. Biotechnology Letters, 44(11), 1337–1346. 10.1007/s10529-022-03300-8

12. Zhang, J., Xue, J., Luo, N., Chen, F., Chen, B., & Zhao, Y. (2023). Microwell array chip-based single-cell analysis. Lab on a Chip, 23(5), 1066–1079. 10.1039/d2lc00667g

13. Lu, X., Pritko, D. J., Abravanel, M. E., Huggins, J. R., Ogunleye, O., Biswas, T., Ashy, K. C., Woods, S. K., Livingston, M. W. T., Blenner, M. A., & Birtwistle, M. R. (2025). Genetically Encoded Fluorescence Barcodes Allow for Single-Cell Analysis via Spectral Flow Cytometry. ACS Synthetic Biology. 10.1021/acssynbio.4c00807

14. Mosadeghi, R., Foyt, D., Sharp, L., Taylor, C., Tay, N., Oberlin, S., Fan, J., Bourke, S., Kattah, M., Huang, B., & McManus, M. T. (2025). RainBar: Optical Barcoding for Pooled Live-Cell Imaging with Single-Cell Resolution. bioRxiv, 2025.11.04.686676. 10.1101/2025.11.04.686676

15. Kawasaki, F., Mimori, T., Mori, Y., Aburatani, H., Yachie, N., Sato, I., & Ota, S. (2023). Computational Design of Synthetic Optical Barcodes in Microdroplets. Advanced Optical Materials, 2302564. 10.1002/adom.202302564

16. Sart, S., Ronteix, G., Jain, S., Amselem, G., & Baroud, C. N. (2022). Cell Culture in Microfluidic Droplets. Chemical Reviews, 122(7), 7061–7096. 10.1021/acs.chemrev.1c00666

17. Shapiro, H. M. (2005). Practical Flow Cytometry (4th ed.). John Wiley & Sons.

18. Perfetto, S. P., Chattopadhyay, P. K., & Roederer, M. (2004). Seventeen-colour flow cytometry: Unravelling the immune system. Nature Reviews Immunology, 4(8), 648–655. 10.1038/nri1416

19. Han, Y., Gu, Y., Zhang, A. C., & Lo, Y.-H. (2016). Review: Imaging technologies for flow cytometry. Lab on a Chip, 16(24), 4639–4647. 10.1039/C6LC01063F

20. Ugawa, M., & Ota, S. (2024). Recent Technologies on 2D and 3D Imaging Flow Cytometry. Cells, 13(24), 2073. 10.3390/cells13242073

21. Martino, N., Kwok, S. J. J., Liapis, A. C., Forward, S., Jang, H., Kim, H.-M., Wu, S. J., Wu, J., Dannenberg, P. H., Jang, S.-J., Lee, Y.-H., & Yun, S.-H. (2019). Wavelength-encoded laser particles for massively multiplexed cell tagging. Nature Photonics, 13(10), 720–727. 10.1038/s41566-019-0489-0

22. Kwok, S. J. J., Forward, S., Fahlberg, M. D., Assita, E. R., Cosgriff, S., Lee, S. H., Abbott, G. R., Zhu, H., Minasian, N. H., Vote, A. S., Martino, N., & Yun, S.-H. (2024). High-dimensional multi-pass flow cytometry via spectrally encoded cellular barcoding. Nature Biomedical Engineering, 8(3), 310–324. 10.1038/s41551-023-01144-9

23. Pregibon, D. C., Toner, M., & Doyle, P. S. (2007). Multifunctional Encoded Particles for High-Throughput Biomolecule Analysis. Science, 315(5817), 1393–1396. 10.1126/science.1134929

24. Utech, S., Prodanovic, R., Mao, A. S., Ostafe, R., Mooney, D. J., & Weitz, D. A. (2015). Microfluidic Generation of Monodisperse, Structurally Homogeneous Alginate Microgels for Cell Encapsulation and 3D Cell Culture. Advanced Healthcare Materials, 4(11), 1628–1633. 10.1002/adhm.201500021

25. Shao, F., Yu, L., Zhang, Y., An, C., Zhang, H., Zhang, Y., Xiong, Y., & Wang, H. (2020). Microfluidic Encapsulation of Single Cells by Alginate Microgels Using a Trigger-Gellified Strategy. Frontiers in Bioengineering and Biotechnology, 8, 583065. 10.3389/fbioe.2020.583065

26. Andersen, T., Auk-Emblem, P., & Dornish, M. (2015). 3D Cell Culture in Alginate Hydrogels. Microarrays, 4(2), 133–161. 10.3390/microarrays4020133

27. Kang, S.-M., Lee, J.-H., Huh, Y. S., & Takayama, S. (2021). Alginate Microencapsulation for Three-Dimensional In Vitro Cell Culture. ACS Biomaterials Science & Engineering, 7(7), 2864–2879. 10.1021/acsbiomaterials.0c00457

28. Yew, P. Y. M., Chee, P. L., Lin, Q., Owh, C., Li, J., Dou, Q. Q., Loh, X. J., Kai, D., & Zhang, Y. (2024). Hydrogel for light delivery in biomedical applications. Bioactive Materials, 37, 407–423. 10.1016/j.bioactmat.2024.03.031

29. Liu, K., Ding, H.-J., Liu, J., Chen, Y., & Zhao, X.-Z. (2006). Shape-Controlled Production of Biodegradable Calcium Alginate Gel Microparticles Using a Novel Microfluidic Device. Langmuir, 22(22), 9453–9457. 10.1021/la061729+

30. Ugawa, M., & Ota, S. (2022). High-Throughput Parallel Optofluidic 3D-Imaging Flow Cytometry. Small Science, 2, 2100126. 10.1002/smsc.202100126

31. Yamashita, M., Tamamitsu, M., Kirisako, H., Goda, Y., Chen, X., Hattori, K., & Ota, S. (2024). High-Throughput 3D Imaging Flow Cytometry of Suspended Adherent 3D Cell Cultures. Small Methods, 8(8), 2301318. 10.1002/smtd.202301318

32. Marini, M. A., Evans, W. J., & Berger, R. L. (1986). The determination of binding constants with a differential thermal and potentiometric titration apparatus. II. EDTA, EGTA and calcium. Journal of Biochemical and Biophysical Methods, 12(3), 135–146. 10.1016/0165-022X(86)90028-X

33. Collins, D. J., Neild, A., deMello, A., Liu, A.-Q., & Ai, Y. (2015). The Poisson distribution and beyond: Methods for microfluidic droplet production and single cell encapsulation. Lab on a Chip, 15(17), 3439–3459. 10.1039/c5lc00614g

34. Dowski, E. R., & Cathey, W. T. (1995). Extended depth of field through wave-front coding. Applied Optics, 34(11), 1859. 10.1364/AO.34.001859

35. Ortyn, W. E., Perry, D. J., Venkatachalam, V., Liang, L., Hall, B. E., Frost, K., & Basiji, D. A. (2007). Extended depth of field imaging for high speed cell analysis. Cytometry Part A, 71A(4), 215–231. 10.1002/cyto.a.20370

36. Dunsby, C. (2008). Optically sectioned imaging by oblique plane microscopy. Optics Express, 16(25), 20306–20306. 10.1364/OE.16.020306

37. Li, T., Ota, S., Kim, J., Wong, Z. J., Wang, Y., Yin, X., & Zhang, X. (2015). Axial Plane Optical Microscopy. Scientific Reports, 4(1), 7253–7253. 10.1038/srep07253

