## Supplementary Information for "Randomized Spatial Barcoding for Time-Lapse Flow Cytometry"

**Estimate of spatial barcode configuration**

The number of spatial configurations that beads encapsulated in a hydrogel droplet can be estimated based on the number of positions each bead can take. In the case of a spherical droplet of 50 μm in diameter and beads of 4.5 μm in diameter, the volume that the center of the first bead can position will be

$$\frac{4}{3}\pi\left(\frac{50-5}{2}\right)^3 = 47713\ \mu\text{m}^3$$

Assuming a resolution of 2 μm in all dimensions, the number of resolvable positions that the bead can take is $47713 \div 2^3 = 5964$. Then, because the center of the next bead must be at least 5 μm away from the previous bead, a 5-μm-radius sphere around the center of the previous bead becomes excluded from the volume. Therefore, the volume that the center of the next bead can take can be calculated as

$$47713 - \frac{4}{3}\pi \times 5^3 = 47189\ \mu\text{m}^3$$

and the number of resolvable position that the next bead can take will be $47189 \div 2^3 = 5899$. For $n$ number of beads, the total number of spatial configurations can be estimated as

$$\frac{\prod_{k=0}^{n-1}\left\{\frac{4}{3}\pi\left(\left(\frac{50-5}{2}\right)^3 - 5^3 \times k\right) \times \frac{1}{2^3}\right\}}{n! \times 4\pi\left(\frac{50-5}{2}\right)^2 \times \frac{1}{2^2}}$$

where $n!$ accounts for the beads being identical, and the last two terms in the divisor accounts for the rotational symmetry. For $n = 3$, this becomes $2.1 \times 10^7$, and for $n = 4$, this becomes $3.1 \times 10^{10}$. Although this overestimates (i) the volume that the beads exclude at the edges of the droplet or when the beads are close to each other and (ii) the number of equivalent configurations based on rotational symmetry, it provides a lower bound estimate to the possible number of configurations.

For the RASPBerry droplets, we used beads with a diameter of 4.5 μm and a hydrogel droplet of an obround shape of 40 μm in width and 60 μm in length. Assuming that all beads are in the same plane, if calculated based on the area in a similar way to above, the total number of spatial configurations for n number of beads can be estimated as

$$\frac{\prod_{k=0}^{n-1}\left\{\left(\pi\left(\frac{40-4.5}{2}\right)^2 + (60-40) \times (40-4.5) - \pi\, 4.5^2 \times k\right) \times \frac{1}{2^2}\right\}}{n! \times 4}$$

where the last term in the divisor accounts for the symmetry of the obround shape. For $n = 3$, 4, and 5, this becomes $2.8 \times 10^6$, $2.7 \times 10^8$, and $1.9 \times 10^{10}$, respectively.

**Estimate of the maximum number of droplets that can be imaged**

The field of view of the line-scan microscopy system is 1,500 μm. The NA of the objective is 0.3 (Supplementary Figure 2), and the resolution is about 1 μm. Using a sampling rate based on the Rayleigh criterion, a 0.5 μm/frame sampling in the flow direction is optimal. Because each channel runs at 2,000/3 frames per second, the flow velocity becomes 333 μm/s. Multiplying this by the field of view of the line image, the imaged area per second is 500,000 $\mu\text{m}^2$/s. Assuming that the geometry of the RASPBerry droplets can be simplified as a 40 × 60 μm rectangle, the maximum droplets that can be imaged per second is roughly 200. If a 1 μm/frame sampling is used, the flow velocity will be 667 μm/s and the droplets that can be imaged per second will be 400.

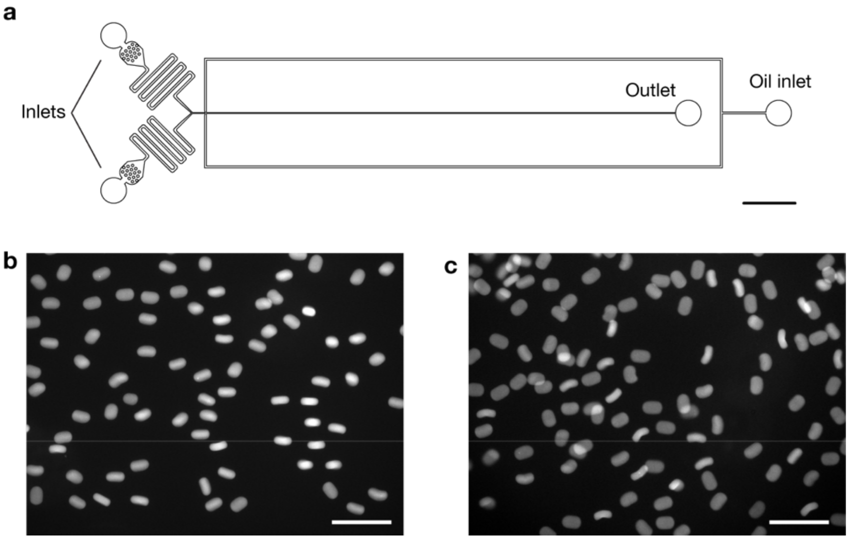


**Supplementary Figure 1.** Fabrication of RASPBerry droplets. (a) Channel design of microfluidic device for fabricating the RASPBerry droplets. Scale bar = 2 mm (b,c) RASPBerry droplets after extraction from oil phase. When the length of the droplets are 60 µm or less the droplets are rarely bent (b), but when the length becomes larger than 65 µm bent droplets become apparent (c). Scale bars = 200 µm

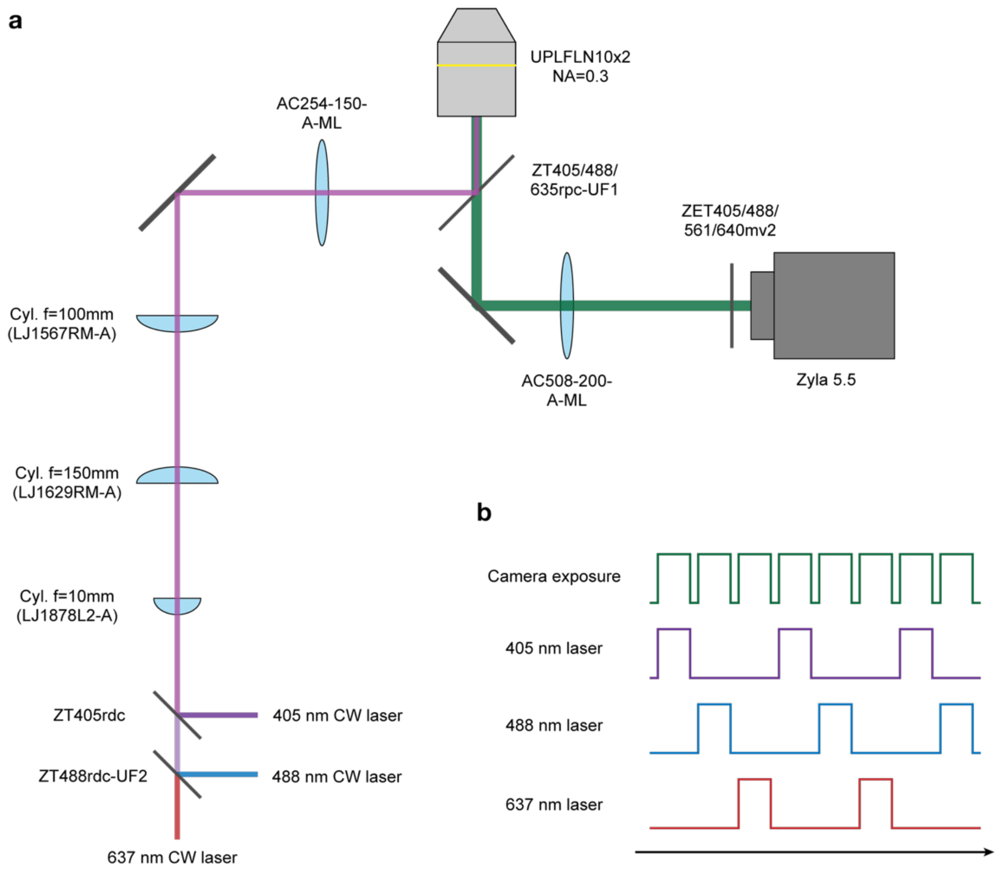


**Supplementary Figure 2.** Optical setup for line-scan microscopy system. (a) Optical schematic of the system. (b) Electronic trigger signals used for operating the lasers and the camera. A 2 kHz signal was used to trigger the acquisition on the camera. The same signal was split into alternating signals using an Arduino microcontroller which were directed to each of the lasers.

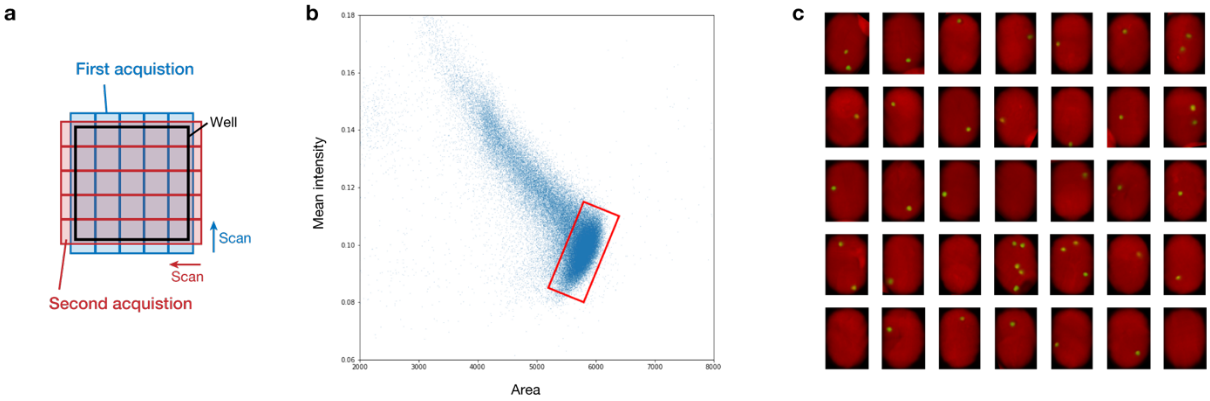


**Supplementary Figure 3.** Imaging of the RASPBerry droplets in a well. (a) Schematic of acquisition for first and second acquisitions. The well was rotated 90 degrees to scan from different directions. (b) Scatter plot of area and mean fluorescence intensity from the segmented RASPBerry droplets from the acquired image. The region within the red rectangle was used to extract the droplets with the flat face parallel to the image plane. (c) 35 mismatched droplets. Most of the droplets have two or fewer beads inside which was not sufficient information for the pattern matching algorithm to detect the same droplet.
